# Community-led research discovers links between elusive symptoms and clinical tests

**DOI:** 10.1101/139014

**Authors:** Irene S. Gabashvili

**Author notes:** Competing interest statement: The author has declared that no competing interests exist.

## Abstract

Human breath and body odors have been used for diagnosis of serious and life-threatening conditions since the dawn of medical practice. More recently, it has been recognized that malodors without accompanying physical symptoms could be a sign of psychologically but not physically debilitating disorders such as Trimethylaminuria (TMAU). Self-reported intermittent odors without apparent cause, are, however, still treated with suspicion by medical professionals. Most cases of socially-disabling idiopathic malodor remain undiagnosed and there are no guidelines for diagnostic tests nor treatment options that extend beyond TMAU. Internationally-recruited volunteers with undiagnosed body odor and halitosis enrolled to participate in our study, registered as NCT02692495 at clinicaltrials.gov. Each volunteer underwent several blood and urine tests conducted by Biolab Medical Unit, a medical referral laboratory in London, specializing in nutritional and environmental medicine. Intestinal permeability measurements were strikingly different for subjects that named the nose/mouth as the malodor source(s) versus other, often unidentified, body regions. Furthermore, metabolite levels in blood and urine allowed matching of participants by dietary sensitivities and the type of odor reported, emphasizing the potential of harnessing patients’ olfactory observations. In discussing the anecdotal “People are Allergic to Me” condition (PATM), we show how it fits into the picture.

## 1. Introduction

To allocate limited resources for medical research, diseases are prioritized. Near the bottom of the prioritized list is trimethylaminuria - an inborn error of choline metabolism that leads to the excessive excretion of foul-smelling trimethylamine (TMA) in the sweat and breath. TMAU sufferers may be physically healthier than the average population, developing gut microbiota that potentially protects them from cardiovascular disease [1]. The same could be said for other sufferers of idiopathic malodor – for example, children with bad breath but good teeth [2] or underarm odor sufferers that harbor less acne bacteria [3]. Yet, psychological, social, and economic implications of uncontrollable odors can have a devastating impact on health.

The rise of online health communities enabled sufferers of rare diseases to communicate with their peers and exchange first-hand knowledge of their conditions. But the only existing diagnostic test for trimethylaminuria is useful for less than 30% of sufferers with the most severe cases [4]. Not having a diagnosis takes an enormous emotional toll. To engage researchers in projects addressing community needs, in 2009, a few sufferers of socially-disabling malodor identified non-standard diagnostic tests that typically yielded positive results, and initiated a study to find if there were clinical similarities with others like them. In this paper, we present results of investigations conducted in 2009-2013 and the most common patterns observed.

## 2. Materials and Methods

### 2.1 Participants

For this Internet-based self-selection trial, 16 volunteers, 11 men and 5 women, aged 22 - 55 years, filled out questionnaires and carried out blood, urine, and breath tests at Biolab’s location in London. Study participants had experienced recurrent episodes of undesirable and uncontrollable odor originating in the oral or nasal cavity (4 subjects) or other areas of the body (12 subjects). The first appearance of persistent odor symptoms (as noticed by others) among the participants was nearly always in young adulthood and about 10 years earlier in male than female participants. The mean duration of symptoms was 9 years. All study volunteers had seen a General Practitioner (GP) and medical specialists about their odor symptoms but had not been given a definite diagnosis. Neither participant of our study had been successfully treated. Six participants underwent trimethylamine challenge, but only one tested positive on second attempt and was suggested the diagnosis of secondary trimethylaminuria [5] – acquired condition that occurs when the liver FMO3 enzyme is either overwhelmed or under-active for dietary, gastrointestinal metabolism, hormonal or other reasons. We note that this was the only volunteer that had malodor experience for only a year at the time of testing, while all others experienced the symptoms from 3 to over 20 years.

### 2.2 Procedures and Data analysis

Biolab Medical Unit, www.biolab.co.uk carried out all diagnostic tests including Gut Permeability Profile, Gut Fermentation Profile, Hydrogen Breath test, functional blood B vitamin profile, plasma D-lactate, and urine Indicans. These tests are described at http://www.biolab.co.uk/index.php/cmsid_biolab_tests, with related peer-reviewed scientific references, physiology, procedures, and factors that may affect the test results. We have made anonymized data partially publicly available [6]. MeBO Research IRB served as an independent ethics committee to protect the privacy and confidentiality of data concerning the research subjects. A set of R packages and Excel macros was developed for statistical analysis. Aurametrix software was used to analyze dietary patterns.

## 3. Results and Discussion

### 3.1 Intestinal permeability

“Leaky Gut Syndrome”, diagnosis coined by environmental medicine practitioners in the 1970s, to describe increased intestinal permeability, has been the subject of speculation as related to malodor. Mainstream science does not recognize it as a distinct medical condition, but agrees that intestinal epithelium is an important target for disease prevention [7]. Dysfunction of the gastrointestinal barrier – normally open enough to allow absorption of nutrients while tight enough to protect against toxins, antigens, and microbes - may predispose to food allergy, inflammatory bowel diseases, irritable bowel syndrome, Parkinson’s, kidney and liver diseases, diabetes, and celiac disease, often associated with unpleasant odors. Researchers speculate that the gastrointestinal tissue damage increasing intestinal release of choline and lecithin, may be the cause of transient TMAU [8]. A simple procedure to assess intestinal permeability involves the oral administration of nonmetabolizable substances and the subsequent measurement of these substances in the urine. A variety of marker probes can be utilized such as lactulose (molecular weight: 342), mannitol (182), rhamnose (164), cellobiose (342), polyethylene glycol 400 (PEG 400, a mixture of molecules of different sizes with molecular weight usually ranging from 242 to 400 Dalton), PEG 1000 (634-1338), 51Cr-EDTA (341) and 99mTc-DTPA (487). The Biolab gut permeability profile used a low dose of PEG (3 g.) measuring all urine passed for the following six hours at eleven different molecular weights (PEG198, PEG242, PEG286, PEG330, PEG374, PEG418, PEG462, PEG506, PEG550, PEG594, PEG638, extracted by ion exchange chromatography), to establish the quantity of each absorbed through the gut wall. This test is particularly suitable to assess the whole-gut permeability.

Fig. 1 shows results of intestinal permeability tests performed between 2009 and 2011. The profiles clearly divide the participants into two subgroups based on the source of malodor. The “Halitosis” subgroup (breath odors emanating from nose/mouth) has lower intestinal permeability than the “Body Odor” subgroup (odors associated with body regions such as scalp, groin, rectum, or armpits). The mysterious PATM condition - coined by one of the sufferers as an acronym of “People Are Allergic to Me”, in 2007 – aligns with the “Body odor” group. Evidence of this syndrome remains anecdotal, despite thousands of sufferers sharing similar life stories on multiple online support groups. PATM is defined as a condition causing people near the sufferer exhibit sensitivity reactions such as rubbing their noses and eyes, sniffing, sneezing, coughing, and clearing their throat.

**Figure 1.**
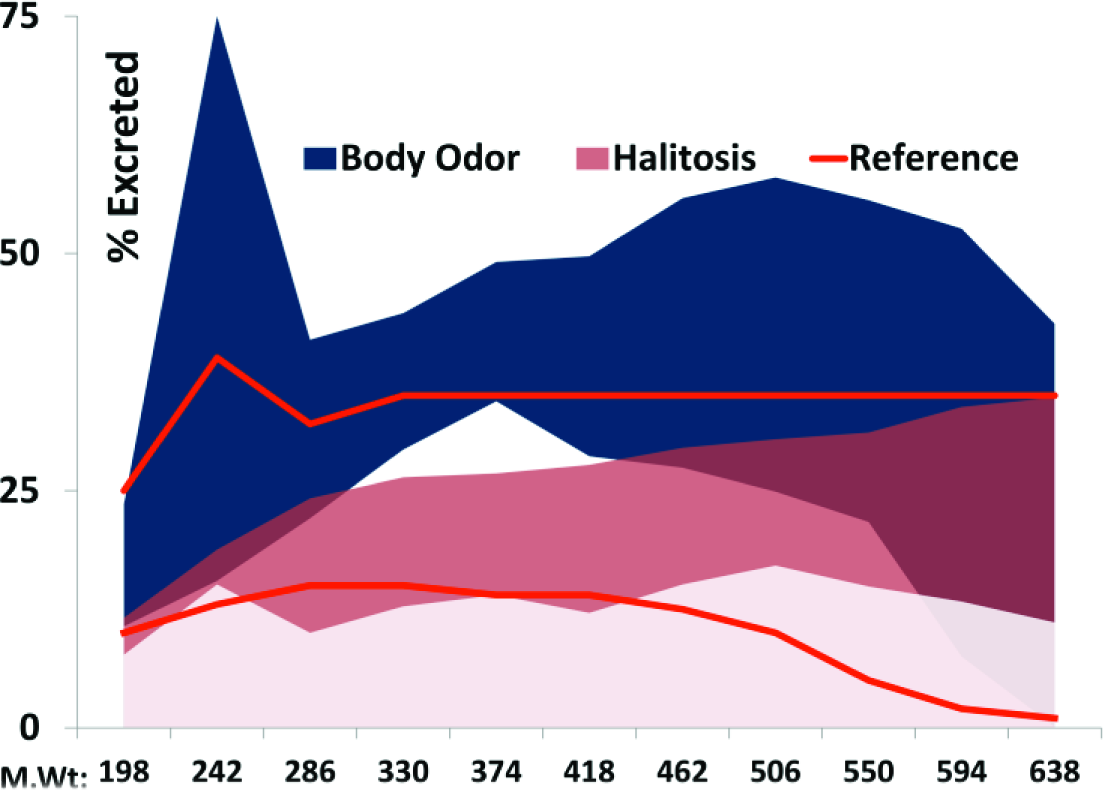
Intestinal permeability profiles grouped by the source of odor reported. Y axes shows percent of PEG (from molecular weight 198 to molecular weight 638) excreted in urine 6 hours after the oral intake.

The “Body odor” profiles appear to lean towards increased permeability and hence “leakage” of microbes and microbial products from the gut into the rest of the body. Gut permeability profiles taken between 2012 and 2013, after recalibration of reference standards by Biolab, while maintained significant differences between the two groups, appeared to be shifted downwards: “Body odor” subgroup was within the reference range, while the “Halitosis” subgroup displayed decreased permeability and malabsorption of essential nutrients. The observation of decreased intestinal permeability in the “Halitosis” group somewhat resembles known cases of mouth odor caused by partial duodenal obstruction [9], although neither of our volunteers have suffered from this or any other significant health problems.

Intestinal permeability data does not confirm nor rule out “leaky gut” (hyperpermeability) and “tight gut” (malabsorption) as causes of malodors, but demonstrates statistically significant differences in intestinal permeability between participants reporting their breath (nose and/or mouth as the malodor source) versus other body regions. The significant differences (*P* < .05) were present between PEG 400 (11.2 Å) and PEG 300 (9.6 Å), approximately where gap junction channels formed by connexin 32 (C×32) have a size cut-off.

### 3.2 Odor indicators

Participants of our study underwent a variety of biochemical tests quantifying about two dozen odorous metabolites, by-products of microbial metabolism, in blood and urine. The molecules studied include D-lactate, Indican, primary, secondary, and tertiary alcohols, short chain fatty acids and related substances.

We summarized these laboratory test results in a 16×14 table with participants numbered from 1 to 16 and 14 metabolites that were identified in participant samples outside expected ranges. We replaced actual measured values of metabolites with discrete “buckets” ranging from “−1” (when test results were significantly lower than in healthy participants not complaining of odors) to 1 (values significantly higher than normal). We analyzed these laboratory test results with several statistical techniques known to bring out strong patterns in a dataset. Results of one such procedure, a principal component analysis (PCA), are shown in Fig.2.

**Figure 2.**
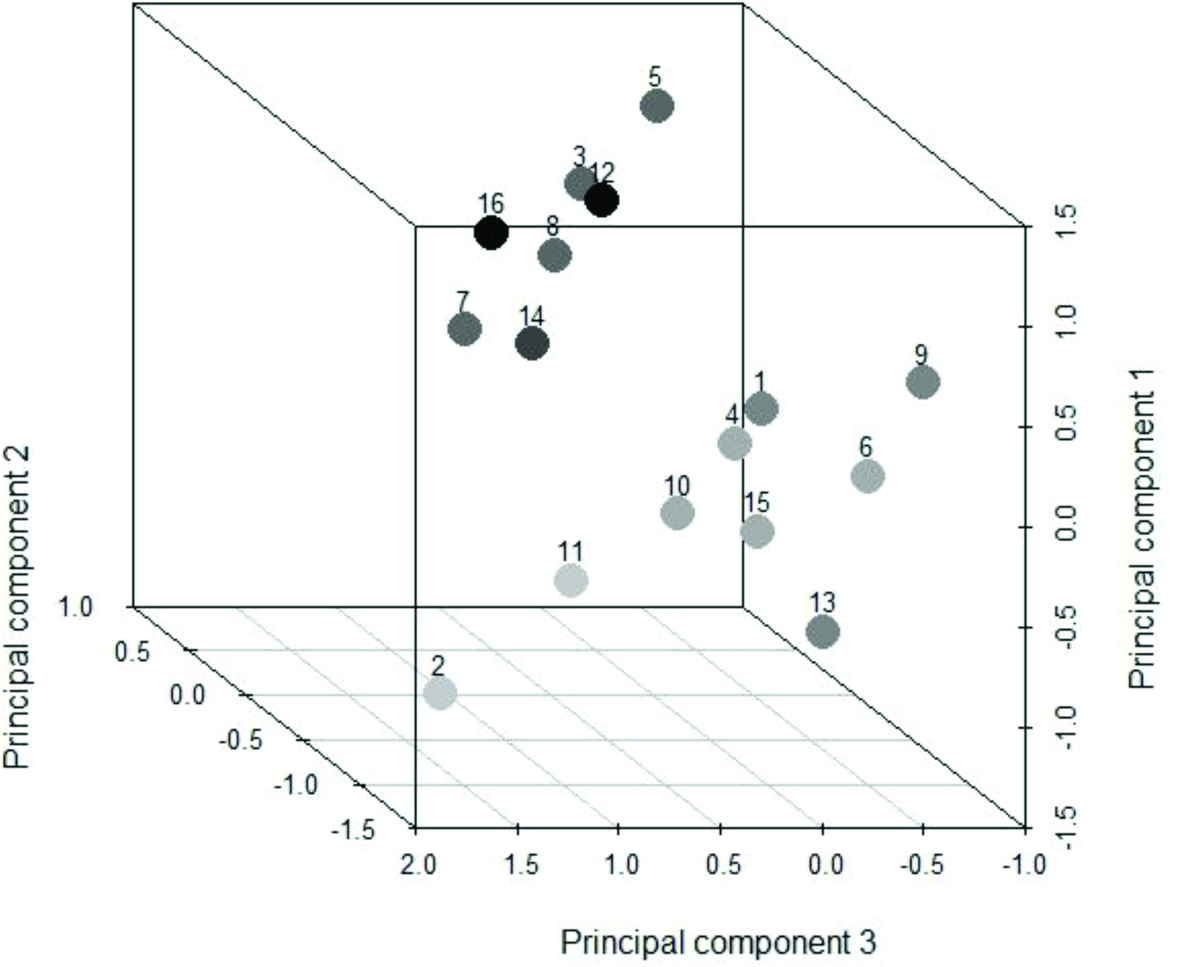
Principal components analysis plot in 3D. The axes show the first three principal components. Each principal component is a linear combination of the original test results indicating over- or under-expression of odorous molecules. Each dot represents a research participant colored per odors reported (see text).

Our main finding was that PCA clustered subjects in groups similar by malodor symptoms reported by the subjects, regardless of the source, age, gender, or other medical history. Another remarkable difference between the two groups was in the participants’ usual consumption of sugar. We will refer to the two main clusters identified as the “sweet” group (darker circles clustered near the top of the cube, in Fig. 2, representing participants with higher sugar intake) and the “sour” group (lighter circles near the bottom, with less sugar in their diets). As actual odors reported by the participants featured over a dozen of keywords, we associated the symptoms with a gamut of gray colors ranging from the lightest – starting from “garbage” odor, followed by “fishy”, “ammonia”, sour acetone (body odors associated with alcoholism), and fecal-diarrheal, followed by generic “fecal odor” - the darkest three circles in the “sour” cluster. The “sweet” group symptom keywords mostly include sweetish sulfurous smells. The odors, from the lightest to the darkest, are “sulfur”, “rotten vegetables”, “rotten eggs”, cheesy/sweaty-fecal, sewage, smoke-like, gasoline, “burning”, followed by those who could not smell themselves and, finally, the “PATM” condition (odors could not be named, but people near the sufferer exhibited increased displeasure, coughing, sneezing, and rubbing their noses). We note that low level exposures to sulfur compounds are known to cause similar issues associated with mild dyspnea.

The axes in Fig. 2 - principal components 1, 2 and 3 - capture a little over 50% of variability in the dataset. Principal component 1 points from sour and acidic odors in detected metabolites (main negative contributors are sour-whiskey-smelling 2-methyl-1-butanol, buttery 2,3-butylene glycol, window-cleaner-like 2-butanol and plastic 2-methyl-1-propanol) towards mild alcohol-like smells such as of 1-propanol. Principal component 3 is somewhat similar – but while its negative values assume more acidic, vinegary, and sour whiskey notes, larger positive values align with other types of sour smells, with sweetish and alcoholic notes. The “sour” subgroup of study participants occupies acidic and sour corner of principal components of the metabolite space, stemming from the lowest values of both PC1 and PC3.

Values of principal component 2 increase as the rancidity and staleness of odors increase – with large positive contributions from rancid-butter-smelling metabolite Butyrate, cheesy Propionate, pungent Acetate and fecal Indican. The “sweet” subgroup is clustered around the areas of highest PC1 (sweet and alcohol-like smells) and PC2.

Another noteworthy observation is that the “sweet” and “sour” groups were significantly different in regards to their sensitivity to probiotic supplements. Three participants from the “sweet” group reported that probiotics improved their odor and only one participant said that the odor worsened after probiotics and vitamins where added to a low choline diet. Neither participant from the “sour” group associated probiotics with improved odor, one participant reported odor getting worse with probiotics but better with plain Greek yogurt, and one participant reported worse odor with antibiotics. Observations about the relationship between odor and meat, dairy, complex and simple carbohydrates or alcohol consumption did not differ between the two groups.

Body levels of B2 vitamin (EGR activation)) were also somewhat correlated with the odor. Most deficiencies were observed in the “sweet” group. Borderline riboflavin deficiency has been also known to cause acute lactic acidosis with symptoms such as sweet-smelling breath.

We note that most existing diagnostic tests on their own were poor separators of self-reported odors and, possibly, its underlying causes. Overexpression of Indican, one of the compounds responsible for the foul smell of feces somewhat correlated with complaints of “fecal-sulfuric” vs other types of fecal odors. Still, the keyword “fecal” was the least descriptive in our study. Plasma D-lactate present at over 150 umol/L could be a good predictor of “garbage” odor, but we don’t have enough data to make definite conclusions. Short bowel syndrome, one of medical conditions associated with overproduction of D-lactic acid, does contribute to unpleasant odors with sour notes, described as “fishy”, “feculent” and “pungent”. Lactic acid molecule, itself, smells musky or mousy, like sour milk – contributing to the “sourness” of the odor.

## 4. Concluding remarks

Our study provides evidence that community research is an important starting point for exploring disorders of unknown origin with elusive symptoms such as intermittent malodor.

Existing diagnostic tests are not sufficient for diagnosing underlying causes for idiopathic malodor, but they help to define subpopulations of patients with elusive symptoms. We found increased levels of blood alcohols (after a glucose challenge test) and urine Indoxyl sulfate in half or more of our participants, but these results had to be combined with additional, lower yielding assays, to better correlate with types of malodor and dietary sensitivities. Opportunistic microorganisms responsible for human malodor and even genetic variations predisposing to higher production of unoxidized trimethylamine are still being defined [10, 11]. Time-consuming and inconvenient for patients trimethylaminuria challenge test is currently the highest-yielding diagnostic procedure with up to 30% positives detected [4] and could be improved if replaced by a metabolome profiling. Intestinal permeability testing provides marginal value for malodor sufferers, but helps to differentiate patients with malodorous breath vs cutaneous respiration.

Diet is known to be associated with body odors. A recent study [12] linked greater fruit and vegetable intake to more pleasant sweet- and medicinal-smelling sweat, and, to a lesser extent, so was fat, meat, egg, and tofu intake, while greater carbohydrate intake was making sweat less pleasant. Our participants, however, found that meat, dairy, fruit and vegetables, probiotics, prebiotics, even low-choline dieting could make them smell worse, proving that a more individualized dietary approach is needed. Low sugar diets, too, do not reduce odors for everyone, as seen from our study: despite other health benefits, sugar reduction didn’t change the unpleasant perception of odors for the “Sour” subgroup of participants, only the types of odors. This is in line with a recent observation that halitosis sufferers eating less sweets had more unpleasantly-smelling volatile compounds in their breath, while children with caries habitually consuming sugar-containing snacks did not have halitosis [2].

Our study represents an important first step forward in understanding unaddressed idiopathic malodors. Our findings emphasize the importance of patients’ self-reported observations and shows the need for comprehensive metabolomic investigation to open new venues in future research toward the successful development of new therapies.

## Acknowledgments

The author received no funding for this work and would like to thank the volunteers of this study and staff of MeBO Research UK, especially Maria de la Torre and Paul Jenkins, for initiating and supporting laboratory testing.

